# Cross-species Comparison of Ultramafic Rock Bio-accelerated Weathering Performance

**DOI:** 10.1101/2024.11.19.624384

**Authors:** Luke Plante, Jacob D. Klug, Joseph Lee, Adrian Hornby, James Adair, Sabrina Marecos, Matthew C. Reid, Esteban Gazel, Buz Barstow

## Abstract

Carbon mineralization is a natural process that sequesters atmospheric CO_2_ by reacting it with cations (*e.g.*, Mg^2+^, Ca^2+^) released by weathering of rocks to form solid carbonates. However, this process is too slow to capture the excess CO_2_ resulting from anthropogenic emissions in time to prevent significant warming of the atmosphere^1–3^. Additionally, accelerating carbon mineralization process by chemical or mechanical methods has proven prohibitively expensive to date^4^. Microbial rock-dissolution processes, including acidolysis, redoxolysis, and complexolysis, have the potential to accelerate weathering with low energy input^5^. However, there is no industrially useful microbe capable of dissolving ultramafic (rich in ferromagnesian minerals) rocks, suggesting that one will need to be discovered or built with synthetic biology. While microbes are known to dissolve ultramafic minerals^6,7^, the performance envelopes for these processes remain uncharacterized, and significant gaps exist in our knowledge of microbe-mineral interaction processes. Here, we make a normalized performance comparison of the dissolution of the ultramafic rock dunite (> 90% olivine ((Mg, Fe)_2_SiO_4_)) by three well-known mineral-dissolving microbes: *Gluconobacter oxydans*^7–9^, *Sphingomonas desiccabilis*^10,11^, and *Penicillium simplicissimum*^12–14^. We show that *G. oxydans* outperformed *P. simplicissimum* and *S. desiccabilis*, producing the most acidic biolixiviant (pH 2.15 when leaching 1% pulp density), and extracting the most magnesium (3,130 mg/L when leaching at 3% pulp density). Additionally, *G. oxydans* co-dissolves nine other metals, eight of which are critical for energy technologies (Cr, Mn, Co, Ni, Cu, and Zn)^15,16^ with a maximum dissolved concentration of 33 mg/L for Ni. While increasing the pulp density of the dunite (solid to liquid ratio) resulted in higher metal dissolution by *G. oxydans* and *S. desiccabilis*, notably pulp densities above 2% inhibited mineral dissolution by *P. simplicissimum*. Our results provide evidence that the gap in performance between *G. oxydans* and the other two microbes increases with pulp density, and thus, *G. oxydans* is best suited for process and genetic engineering to maximize performance and minimize costs and environmental impacts of bio-accelerated weathering. Finally, we propose that *G. oxydans* and *P. simplicissimum* can use cellulosic hydrolysate as a cost-effective substitute for glucose for biolixiviant production.

## Introduction

In 2018, the Intergovernmental Panel on Climate Change (IPCC) estimated that 10 to 20 gigatonnes of CO_2_ must be removed annually from the atmosphere by the end of the century to prevent warming of 1.5 ^°^C above pre-industrial average global temperatures^17^.

Ultramafic rocks (rocks rich in ferromagnesian minerals) provide a unique opportunity for CO_2_ removal. These rocks form in the Earth’s mantle and are far from chemical equilibrium at Earth’s surface and represent a vast reservoir of potential energy for chemical reactions^18^. Natural weathering of these rocks by rainwater liberates carbonate-forming metals like magnesium (Mg^2+^) and calcium (Ca^2+^), ultimately leading to reaction with dissolved inorganic carbon species and carbon mineralization (formation of solid carbonates like magnesite). This process sequesters between 381 million and 1.06 billion tonnes of CO_2_ per year^19,20^. However, this is offset by release of a few hundred megatonnes of CO_2_ each year by oxidation of rock organic carbon^21^. In total, it is estimated that surface-accessible ultramafic rocks have the capacity to sequester between 100 to 100,000 trillion tonnes of CO_2_ as carbonate minerals^22^. This could account for all excess CO_2_ added to the carbon cycle since the start of the Industrial Revolution at least 58 times over^23,24^ (**Note S1**). However, while ultramafic rocks are relatively unstable, their dissolution and subsequent mineralization is still very slow relative to anthropogenic emissions. As a result, these naturally-occurring processes (mineral dissolution and subsequent carbonation) will eventually sequester all excess anthropogenic CO_2_, but are likely to take thousands to hundreds of thousands of years to do it^1–3^ (**Note S2**).

While dissolution of ultramafic rocks by rainwater can be mechanically accelerated^25^, grinding to the size needed to achieve a weathering rate that will meet the IPCC target is prohibitively expensive. The cost of sequestering a tonne of CO_2_ by this method is between at least $60 and $550^4,26^, likely well above the US Department of Energy’s target of $100^27^., and the cost could be even higher if temperature increases above 200 ^°^C were used for process optimization^18^. Thus, to make carbon mineralization viable in terms of rate and cost, natural weathering must be accelerated.

Processed ultramafic mine tailings are a natural starting point for carbon mineralization as they already have a reduced grain size (< 10 mm to 0.1 mm^28^). This reduces estimated CO_2_ sequestration costs to only about $10 per tonne^4^. However, the supply of tailings is limited, and the total sequestration potential is well below the IPCC’s target. It is estimated that there are less than 10 gigatonnes of ultramafic mine tailings on hand^26^, while annual production amounts to approximately 400 million tonnes^29^. If all Mg^2+^ cations released from the tailings react to form carbonates, the stockpiled tailings could sequester 5 gigatonnes of CO_2_, while newly produced tailings could sequester 190 megatonnes of CO_2_ annually (see **Note S3** and **Dataset S1**).

In addition to magnesium, ultramafic rocks also contain other valuable elements including chromium, manganese, cobalt, nickel, and also some copper, and zinc^30^. Extracting these elements may help offset operating costs in large-scale ultramafic leaching and carbon sequestration processes^31^. Many of these metals are also critical in the energy infrastructure development that future generations will demand^15,16^. However, extracting metals (Mg for CO_2_ sequestration, Ni and other metals for energy technologies) from bulk ultramafic rocks is likely prohibitively expensive with current technology. A major component of this processing cost is comminution (*i.e.*, grinding). Furthermore, leaching of metals with inorganic acids also raises significant concerns about environmental impacts^32^. Thus, engineering a way to accelerate bulk ultramafic rock dissolution and reactions with environmentally-friendly solvents, that would otherwise be slow at surface conditions^33,34^, is worthy of investigation.

Bioleaching could offer an alternative, lower-cost, lower-environmental impact way to release metals for carbon mineralization and energy technologies from ultramafic materials. Many microbial species produce mineral-dissolving cocktails called biolixiviants. Biolixiviants dissolve minerals by at least three mechanisms: acid production (acidolysis); chelators that bind to and extract specific elements (complexolysis); adjusting the redox state of the metals (redoxolysis); or a combination of these^8,9,32,35–44^. In some cases, these biolixiviants have been shown to dissolve metals with very high (>80%) efficiency^8,45^. In fact, biomining already makes up 15% of copper production today^46,47^ and 5% of gold production^46^, and efforts to commercialize lithium biomining are underway^48,49^.

While it is known that microbes can interact and potentially bioleach ultramafic rocks^6,7^, further work is required to optimize this leaching. Different microbes known to bioleach other minerals must be compared in their relative performance at ultramafic rock dissolution, the best-performing microbe may be genetically engineered to bioleach with even more efficiency, and process engineering must occur to maximize economic viability. Here, we make a normalized comparison of the ultramafic bioleaching performance of three mineral-dissolving microbes: *Gluconobacter oxydans, Sphingomonas desiccabilis, and Penicillium simplicissimum Gluconobacter oxydans* is a bacterium with significant biomining potential. It has been shown to convert glucose to gluconic acid as a component of its biolixiviant, and it has been used to successfully bioleach rare earth elements (REE)^8,9,35^. Additionally, *G. oxydans*’ metabolic pathways and genetics are relatively well-understood and easy to engineer. Engineering this bacterium to develop mutants capable of leaching ultramafic materials even better than wild-type strains is already well-within the realm of possibility^9^.

*Sphingomonas desiccabilis* has been shown to successfully bioleach minerals including dissolving phosphates^50,51^ and basaltic rocks^10,11^. Additionally, *S. desiccabilis* survives drying to point of desiccation, which can result in significant cost savings in storing and transporting cultures. For these reasons, this microbe was even studied on the International Space Station for REE biomining performance. However, its bioleaching mechanism is unknown as it may not always produce acids in biomining^10,11^. Though *S. desiccabilis* is known to produce extracellular polymeric substances (EPS) that may complex and form uronic acid in some cases, it is also thought to use complexolysis for bioleaching^11,52,53^.

*Penicillium simplicissimum* is a filamentous fungus shown to produce acids capable of effectively dissolving several metals from minerals, including Al, Fe, Mo, and NI^14,54,55^. This fungus has also been shown to produce citric acid, oxalic acid, and gluconic acid in its biolixiviant^14^. The slow initial growth from spores to form a mycelium relative to bacterial growth was anticipated to be the main disadvantage this microbe would have compared to the two bacteria, though previous work suggests that its leachate may indeed have been at its lowest pH with the five days of bioleaching planned for these tests^55^.

Furthermore, while glucose is a favorable bioleaching feedstock for many microbes^7–11,35,55,56^, alternative feedstocks may be required for this bioleaching to occur at scale due to agricultural limitations^57,58^ and cost^59^. Cellulosic hydrolysate is a promising feedstock made from waste materials like corn stover, cardboard^60,61^, non-recyclable paper^61^, date palm clippings^61^, or eucalyptus bark^62^. These feedstocks, currently in development in several laboratories and pilot plants globally, may significantly increase the availability of sustainable microbial feedstocks^61–63^.

Because of the significant differences between *G. oxydans*, *S. desiccabilis*, and *P. simplicissimum*, their relative performance at bioleaching metals from ultramafic material was not known and could not have been confidently predicted prior to this study. Furthermore, it is unclear how the use of cellulosic hydrolysate as a feedstock impacts bioleaching performance. While previous studies have focused on either mixed culture reactors^6^ or *G. oxydans*’ ultramafic bioleaching^7^, a direct comparison of ultramafic bioleaching performance by different species is lacking. Our aim was to identify the best species for this task and to further increase its performance in future work with genetic engineering.

## Results and Discussion

The aim of this study is to compare the relative ultramafic bioleaching performance of *G. oxydans*, *S. desiccabilis*, and *P. simplicissimum* under identical conditions. While these three microbes are all very different, we were able to normalize leaching conditions by using consistent time periods, media, and growing conditions. Bioleaching performance was evaluated at three pulp densities (*i.e.*, the ratio of solid to liquid in the experiment) of the ultramafic rock dunite, primarily using glucose as a feedstock for biolixiviant production (*i.e.*, the microbes were grown to saturation, and then fed glucose to initiate biolixiviant production and bioleaching). Finally, we compared bioleaching performance using cellulosic hydrolysate as a feedstock against glucose.

Under the conditions of these tests, all microbes produced organic acids (**Figures 1** and **2**). Ranked performers at ultramafic bioleaching were *G. oxydans*, *P. simplicissimum*, and *S. desiccabilis* (**Figures 3** to **6**). Leachate pH in almost all experiments was inversely proportional to magnesium and nickel concentrations (**Figure 7**). All mass spectrometry data used in this article is included in **Dataset S2**.

**Figure 1.**
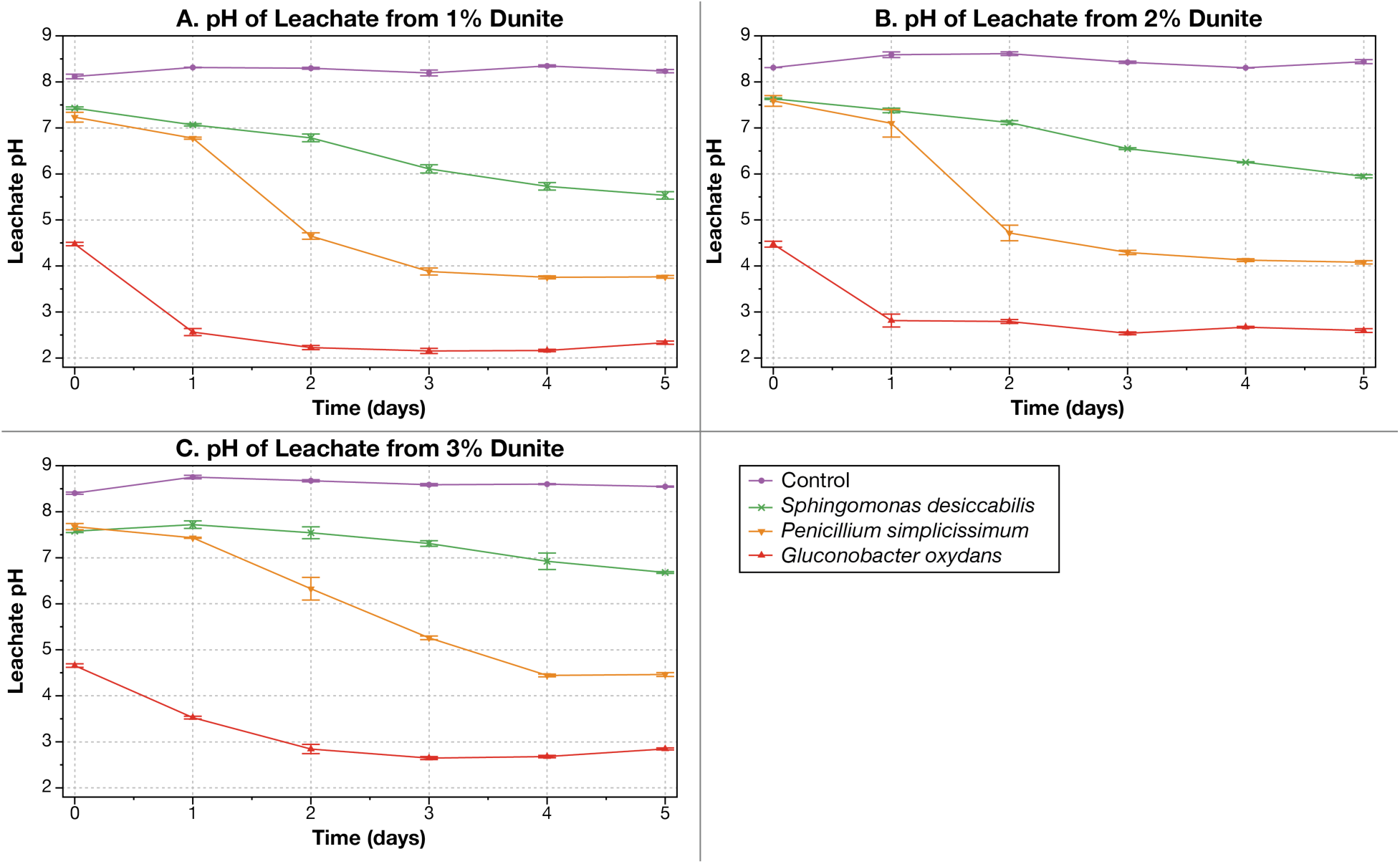
All mineral-dissolving microbes lowered the pH of the leachate from the ultramafic rock dunite through five days. (**A** to **C**) Reduction in pH of dunite leachate at 1, 2, and 3% pulp density (1% pulp density = 1 g per 100 mL of solution). The pH of the leachate produced by *Gluconobacter oxydans* (red triangles) and *Penicillium simplicissimum* (orange inverted triangles) reached their minimum values within 5 days, but the leachate produced by *Sphingomonas desiccabilis* (green crosses) had yet to stabilize. The control was sterile glucose. Raw data for this plot can be found in **Dataset S3**. Error values represent standard deviations. Temperature was maintained at 30 °C throughout the experiment.

**Figure 2.**
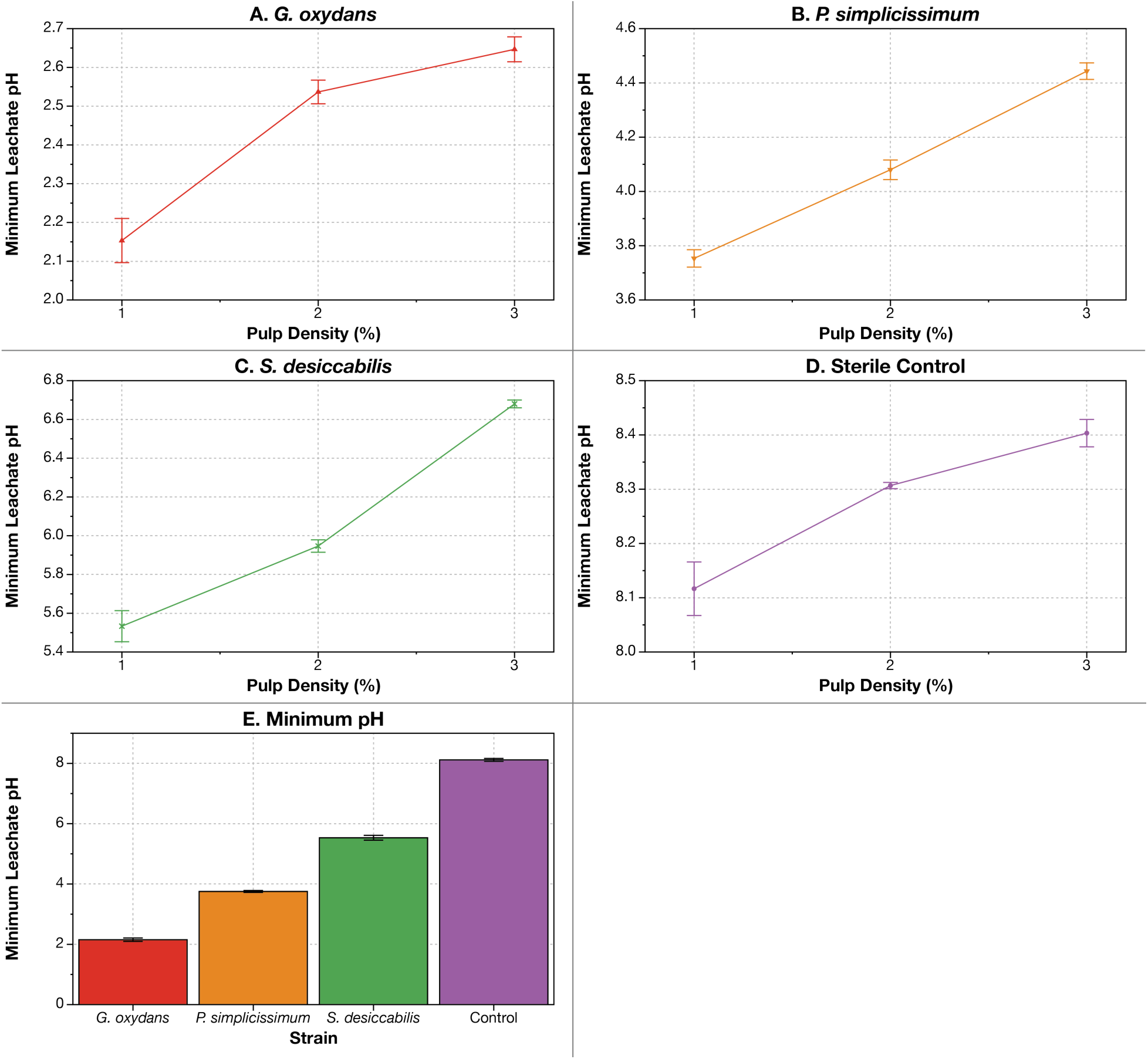
*G. oxydans* produced the most acidic leachate from dunite (A), followed by *P. simplicissimum* (B), and *S. desiccabilis* (C) (summarized in E). (**A** to **D**) pH of leachates produced by all microbes and the sterile glucose controls increased with the pulp density of dunite, likely due to the alkalinity that dunite produces. (**E**) *G. oxydans*’ leachate was most acidic at all pulp densities, with a minimum pH of 2.15 at 1% dunite on day 3. Additional data for this panel is shown in **Table S1**. Data for this plot can be found in **Dataset S3**. Error values represent standard deviations. Temperature was maintained at 30 °C throughout the experiment.

**Figure 3.**
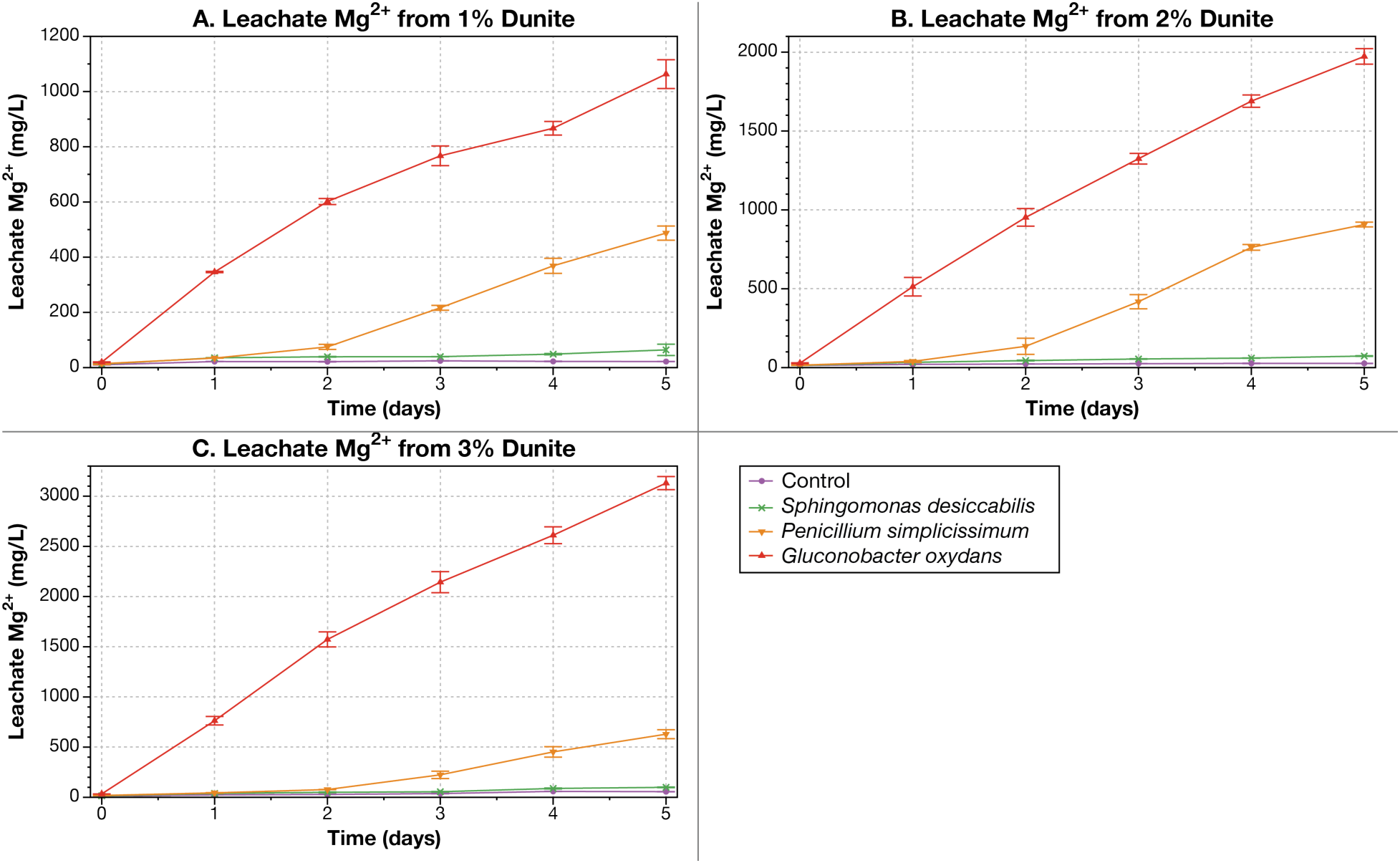
All bioleaching microbes continued to increase the concentration of Mg^2+^ extracted from dunite for all 5 days of bioleaching at 1, 2 and 3% pulp density (A to C). The control was sterile glucose. Raw data for this plot can be found in **Dataset S3**. Error values represent standard deviations. Temperature was maintained at 30 °C throughout the experiment.

**Figure 4.**
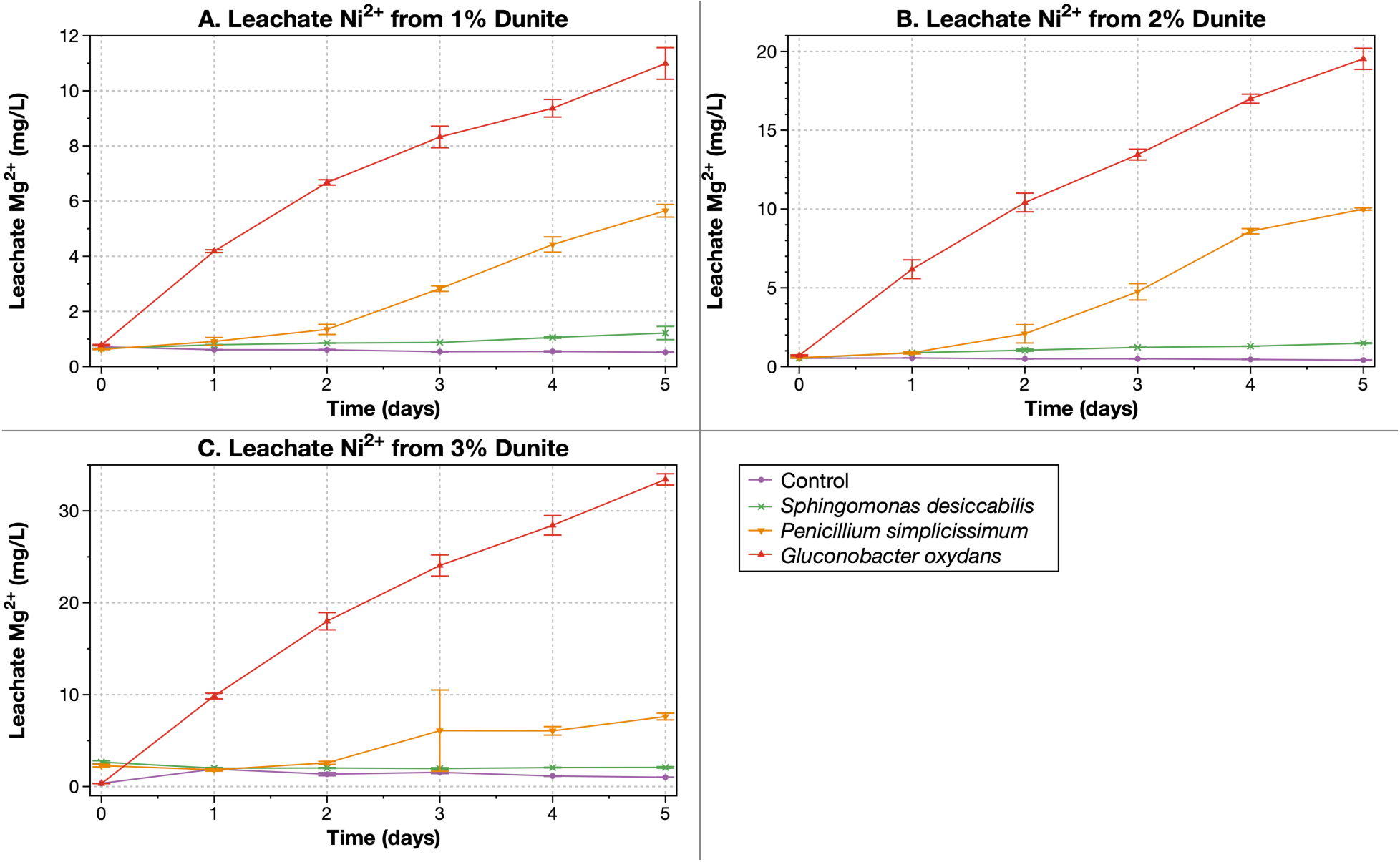
*G. oxydans* and *P. simplicissimum* increased the amount of Ni^2+^ leached from dunite over all 5 days of bioleaching at 1, 2 and 3% pulp density (A to C). Bioleaching of Ni^2+^ by *S. desiccabilis* increased at 1 and 2% pulp density, but was not above the uncertainty in our baseline at 3% pulp density. The control was sterile glucose. Raw data for this plot can be found in **Dataset S3**. Error values represent standard deviations. Temperature was maintained at 30 °C throughout the experiment.

**Figure 5.**
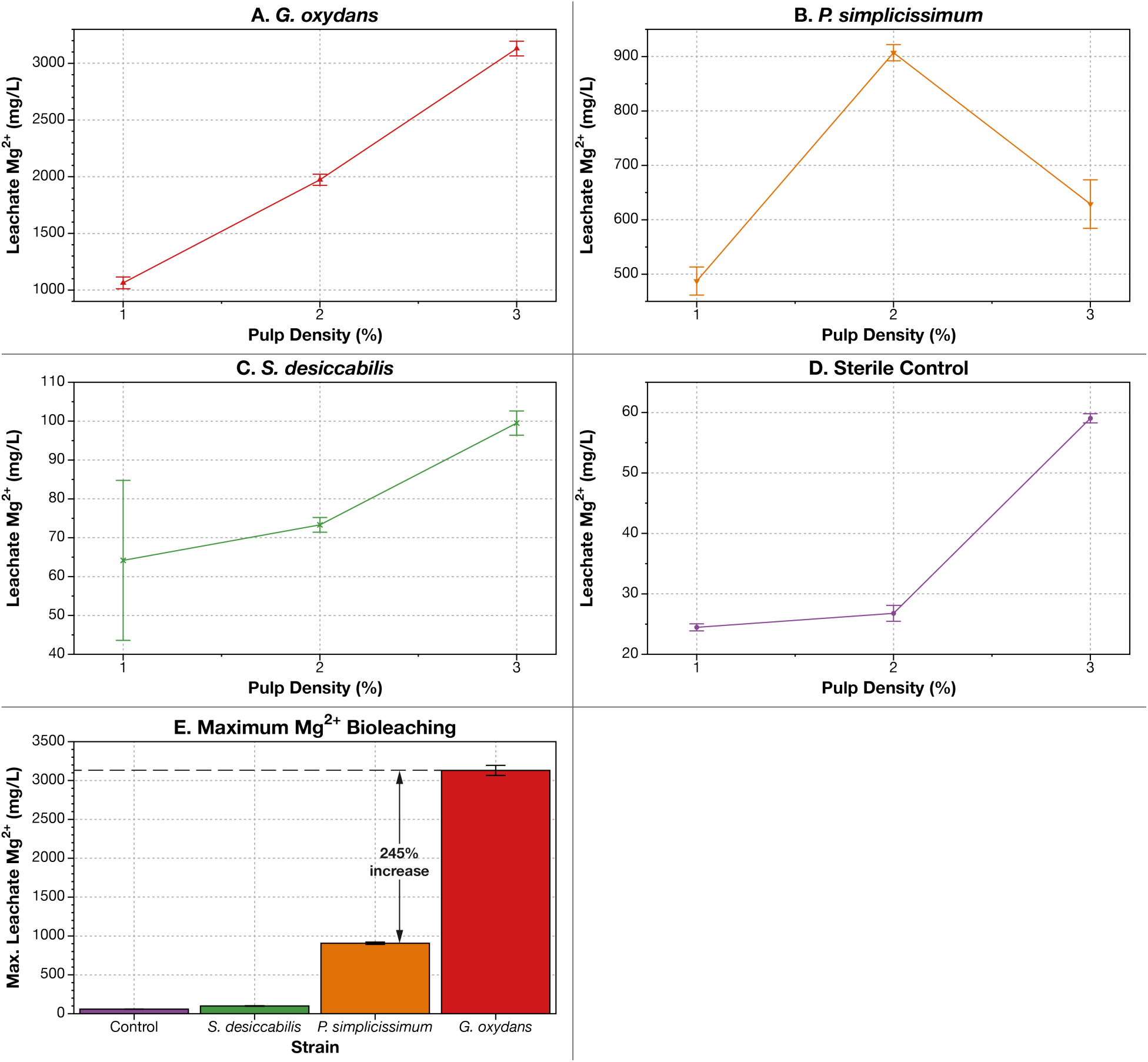
*G. oxydans* produced the highest concentration of Mg^2+^ from dunite (A), followed by *P. simplicissimum* (B), and *S. desiccabilis* (C) (summarized in E). Bioleaching of Mg^2+^ increased roughly proportionally with pulp density for *G. oxydans* (**A**) and *S. desiccabilis* (**C**), but higher concentrations of dunite inhibited *P. simplicissimum* bioleaching (**E**) *G. oxydans* produced a maximum Mg^2+^ concentration of 3,130 mg/L after 5 days of leaching at 3% dunite, over 245% higher than the maximum leaching produced by *P. simplicissimum*. Raw data for this plot can be found in **Dataset S3**. Error values represent standard deviations. Temperature was maintained at 30 °C throughout the experiment.

**Figure 6.**
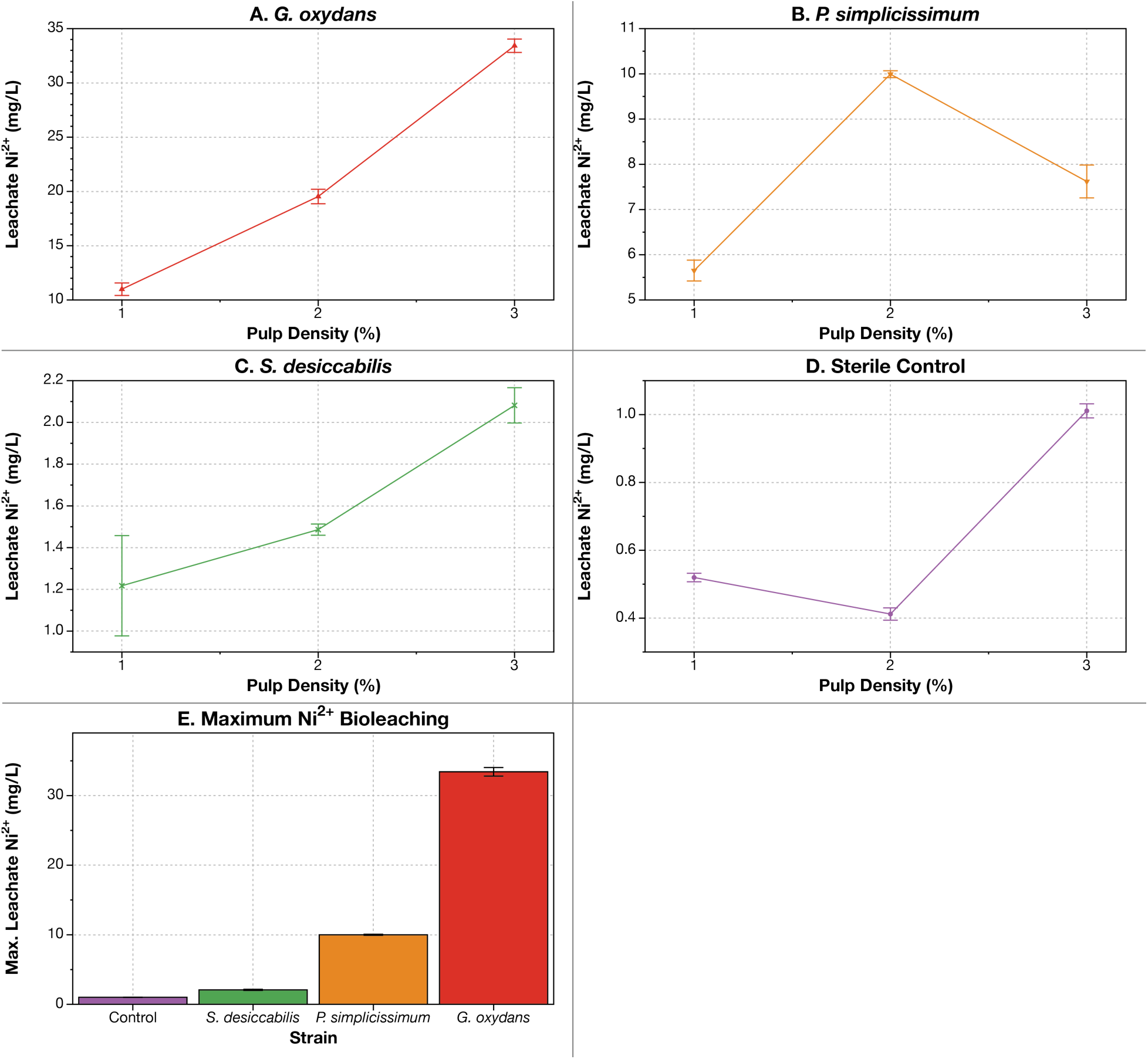
*G. oxydans* produced the highest concentration of Ni^2+^ from dunite (A), followed by *P. simplicissimum* (B), and *S. desiccabilis* (C) (summarized in E). Bioleaching of Ni^2+^ increased roughly proportionally with pulp density for *G. oxydans* (**A**) and *S. desiccabilis* (**C**), but higher concentrations of dunite inhibited *P. simplicissimum* bioleaching (**E**) *G. oxydans* produced a maximum Ni^2+^ concentration of 33 mg/L after 5 days of leaching at 3% dunite, over 234% higher than the maximum leaching produced by *P. simplicissimum*. Raw data for this plot can be found in **Dataset S3**. Error values represent standard deviations. Temperature was maintained at 30 °C throughout the experiment.

**Figure 7.**
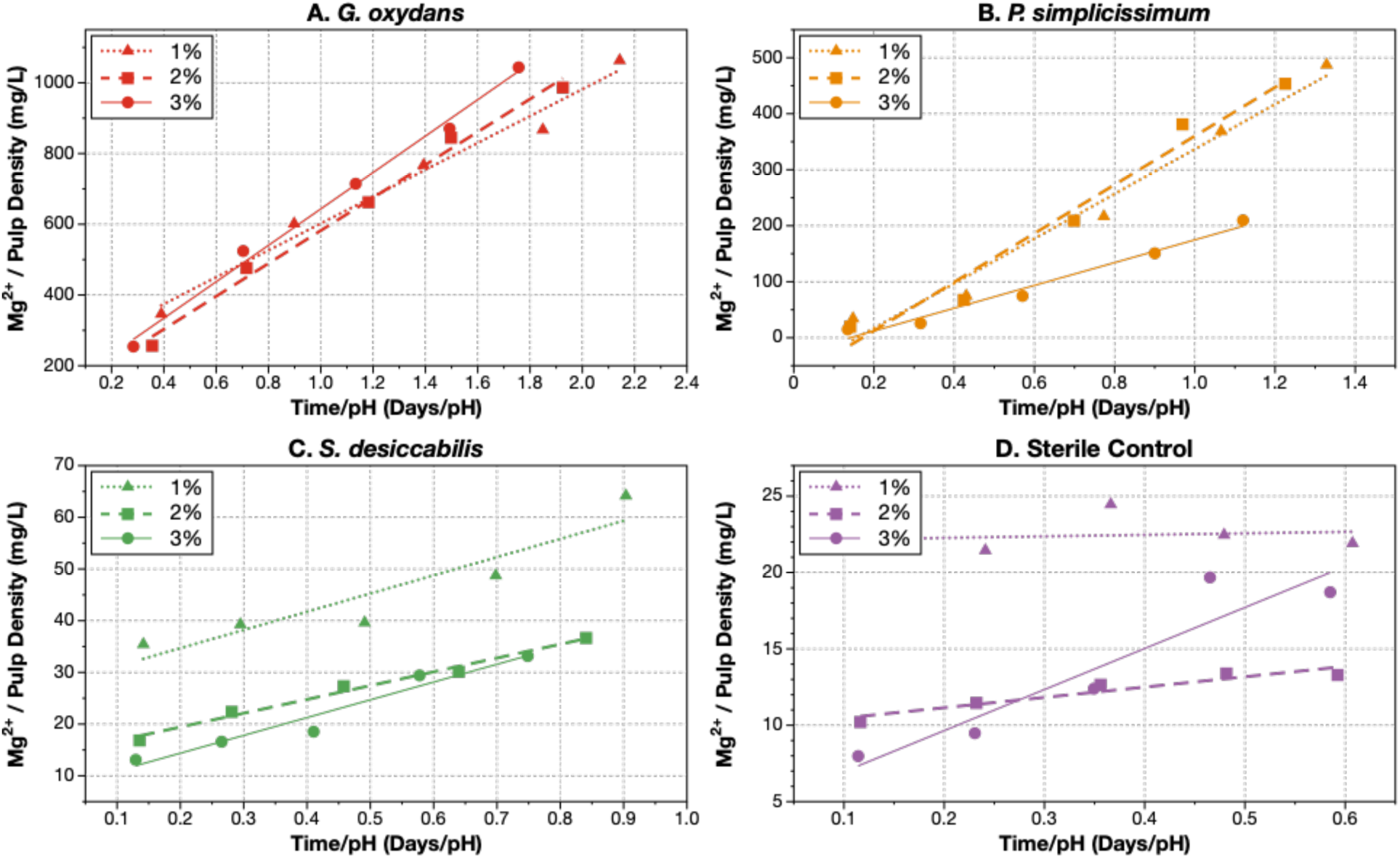
In most cases, bioleaching is inversely proportional to pH and proportional to time. (A) *G. oxydans*; (B) *P. simplicissimum*; (C); *S. desiccabilis*; (D) sterile glucose control.

### *G. oxydans*’ Leachate was Most Acidic by Over 1.5 pH Units

Each mineral-dissolving microbe tested reduced the pH of leachate from dunite through five days of bioleaching at pulp densities of 1, 2, and 3% when using glucose as a feedstock for biolixiviant production (**Figure 1**). Higher pulp densities consistently produced higher leachate pHs (**Figures 2A** to **2D**). This result makes intuitive sense as dunite produces alkalinity in water, so higher pulp densities should increase the leachate pH^64^. Additionally, we hypothesized that higher pulp densities could also prevent very low leachate pHs being achieved by increasing the concentration of potentially toxic metals like iron (II)^65^ and chromium^66^ that may cause stress in some of the bioleaching microbes and a reduction in acid production. These two factors (increased alkalinity and increased toxic metal concentration) could theoretically lead to higher leachate pHs, and higher pHs could result in decreased metal extraction. Conversely, it was also unknown whether the presence of any of these metals in increasing concentrations would stimulate the microbes to bioleach more effectively.

*G. oxydans* produced the most acidic leachate from dunite (2.15 ±0.06, reached after 3 days; **Figures 1A** and **2E**, and **Table S1**). This is comparable to that produced when bioleaching rare earth element-containing materials like retorted phosphor powder, allanite, and monazite^9,35,41^. Under all conditions, the peak acidity of the *G. oxydans* leachate was no less than 1.5 pH units lower than that produced by *P. simplicissimum* or *S. desiccabilis* (**Figure 2E**, and **Table S1**). Minimum pHs for all experiments are summarized in **Table S1**.

The large observed reductions in leachate pH were anticipated for both *G. oxydans* and *P. simplicissimum.* Both organisms are well-known producers of organic acids, and these acids are thought to contribute to leaching^8,9,35,55^. The leachate pHs suggest that their biolixiviants reached their maximum concentrations after 3 to 4 days of bioleaching (**Figure 1** and **Table S1**). On the other hand, *S. desiccabilis* activity was not necessarily expected to decrease the leachate pH. Literature suggests that it may not produce acids as it can bioleach in neutral pH ranges^10^, but it clearly reduces the leachate pH by almost 3 pH units over the sterile control (**Figure 1** and **Table S1**). The constant decrease in *S. desiccabilis* pH requires further investigation to better understand the microbe’s bioleaching mechanisms and to determine its dominant mechanism.

### *G. oxydans* Leached Metals Most Effectively

The acidic biolixiviants produced by all three mineral-dissolving microbes from glucose extracted at least ten metals from dunite: Mg, Fe, Al, Si, Cr, Mn, Co, Ni, Cu, and Zn (**Dataset S2**). In the main text, we have chosen to focus on extraction of magnesium (useful for carbon sequestration) (**Figures 3** and **5**), and nickel (the energy-critical metal with the highest concentration in the leachate) (**Figures 4** and **6**).

Both magnesium (Mg^2+^) and nickel ion (Ni^2+^) concentrations continued to increase (**Figures 3** and **4**) even after leachate pHs reached their floors (**Figure 1**). This suggests that acids were a contributing mechanism for dissolving these metals from dunite, and increased exposure to the acids over time led to increased leaching. Accounting for the exposure time to acids is therefore likely an important factor in developing an optimized industrial ultramafic bioleaching process.

### Higher Pulp Densities Tended to Produce Higher Metal Bioleaching

At any given pulp density, lower leachate pHs consistently produced higher Mg^2+^ and Ni^2+^ concentrations (**Figure 7**). As a result, under the conditions in each of these tests, the top performer in bioleaching Mg^2+^ was *G. oxydans*, followed by *P. simplicissimum* and *S. desiccabilis* (**Figure 5E**). Likewise, *G. oxydans* also produced the highest Ni^2+^ bioleaching, also followed by *P. simplicissimum* and *S. desiccabilis* (**Figure 6E**).

Furthermore, higher pulp densities generally correlated with higher leachate metal concentrations (**Figures 5A** to **D**, and **6A** to **D**). *G. oxydans* produced a maximum Mg^2+^ concentration of 3,130 mg/L after 5 days of bioleaching at 3% pulp density (**Figure 5E**). Similarly, *G. oxydans* produced a maximum Ni^2+^ concentration of 33 mg/L after 5 days of bioleaching at 3% pulp density. Maximum Mg^2+^ and Ni^2+^ concentrations are further summarized in **Table S2**.

### Higher Pulp Density of Dunite Inhibits *P. simplicissimum*’s Bioleaching Performance

While metal concentrations increased with pulp density in leachates produced by *G. oxydans*, *S. desiccabilis*, and even the sterile glucose control (**Figures 5A**, **5C**, **5D, 6A**, **6C**, and **6D**) this was not the case for *P. simplicissimum*. Bioleaching of Mg^2+^ and Ni^2+^ by *P. simplicissimum* peaked at 2% pulp density, before falling at 3% (**Figures 5B** and **6B**). Bioleaching by *P. simplicissimum* was approximately half as effective as that by *G. oxydans* at 1% and 2% pulp density but decreased to less than a quarter at 3% (**Figures 5A** and **6A** versus **5B** and **6B**). At 10% pulp density, *P. simplicissimum* bioleaching of Mg^2+^ using glucose was only about 1/6^th^ as effective as bioleaching by *G. oxydans* using glucose after 15 days (**Figure 9**).

Acid production by *P. simplicissimum* notably lagged at 3% pulp density (**Figure 8A**). While the pH of the leachate produced by *P. simplicissimum* at 1 and 2% dunite reached its minimum value within 2 or 3 days, this stabilization did not occur until day 4 of leaching at 3% pulp density (**Figure 8A**). This time lag explains at least some of the decreased metal bioleaching by *P. simplicissimum* (**Figure 8B**).

**Figure 8.**
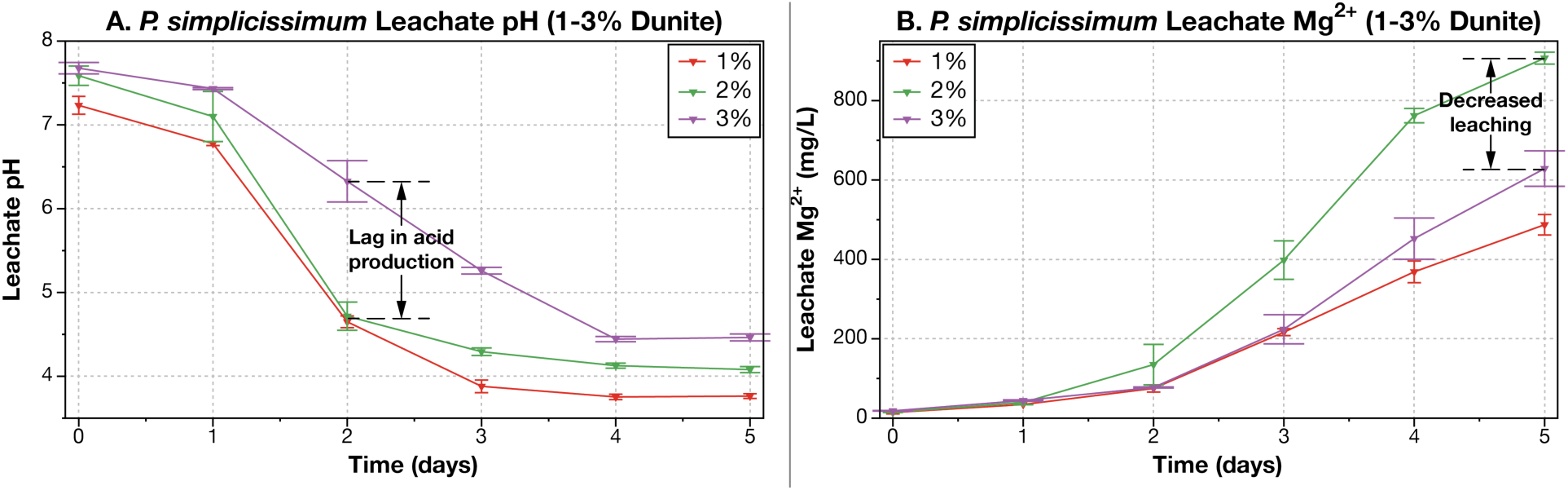
Higher dunite pulp density inhibited bioleaching by *P. simplicissimum* due to a lag in acid production. (**A**) *P. simplicissimum* showed a lag in acid production at 3% dunite pulp density (purple inverted triangles) compared with 1% and 2% dunite. (**B**) Bioleaching at 3% pulp density also lagged behind the leachate magnesium at 2%. This inhibition was not observed in either of the other two species tested. Raw data for this figure can be found in **Dataset S3**. Error values represent standard deviations. Temperature was maintained at 30 °C throughout the experiment.

These results suggest that *P. simplicissimum* is sensitive to some part of dunite’s composition. An element or compound in dunite may inhibit *P. simplicissimum’s* bioleaching performance above a certain pulp density. Notably, *G. oxydans* is far more tolerant to this effect, but not totally immune to it. At 10% pulp density, *G. oxydans* extracts 8,800 ± 160 mg/L of Mg^2+^, less than what would be expected if the approximately linear trend in bioleaching with pulp density in **Figure 5A** had continued (note: a direct comparison is not possible, as bioleaching in **Figure 9** occurred over 15 days, while in **Figure 5** it occurred over 5 days, and temperatures were at room temperature (21 to 23 °C) for testing at 10% pulp density and 30°C for testing at 1%, 2%, and 3% pulp densities).

**Figure 9.**
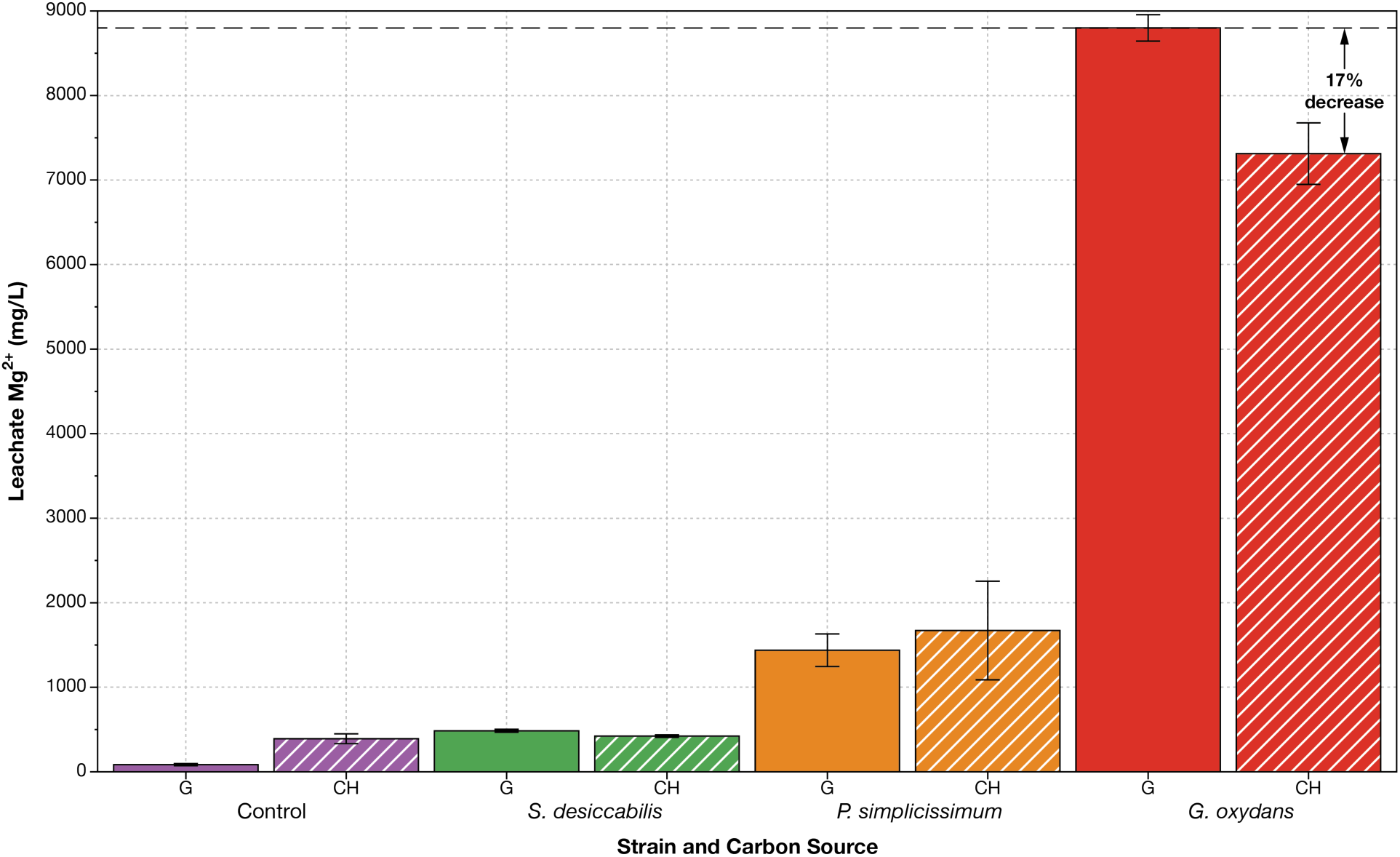
Cellulosic hydrolysate (CH) shows promise as a low-cost, environmentally-friendly alternative to glucose feedstocks for bioleaching (G). *G. oxydans* and *P. simplicissimum* each successfully bioleached magnesium from dunite using cellulosic hydrolysate instead of glucose. Raw data for this plot can be found in **Dataset S3**. Error values represent standard deviations. Cellulosic hydrolysate controls showed a 264% increase in performance compared with glucose controls. Experiment was performed at room temperature (21 to 23 °C).

Further investigation is required to elucidate these inhibiting factors, but this constraint suggests that at least at conditions similar to those used in this test, *G. oxydans* is much better suited for large-scale ultramafic bioleaching than is *P. simplicissimum*.

### Cellulosic Hydrolysate is Viable as an Alternative Feedstock to Glucose

While glucose appears to be effective as a feedstock for bioleaching (especially for *G. oxydans*), its relatively low availability and relatively high cost could seriously limit the scale and cost-effectiveness of biomining ultramafic material, though recent developments show promise for increased availability and decreased cost^67,68^. We tested if cellulosic hydrolysate could serve as an effective substitute for glucose.

Under conditions of 10% pulp density and 15 days of bioleaching, both *G. oxydans* and *P. simplicissimum* effectively bioleached magnesium from dunite using both glucose and cellulosic hydrolysate as feedstocks (**Figure 9**). As we noted earlier, bioleaching by *P. simplicissimum* was only about 1/6^th^ as effective as that by *G. oxydans* using glucose. Notably, bioleaching with by *P. simplicissimum* was not significantly different when using cellulosic hydrolysate or glucose. Furthermore, the use of cellulosic hydrolysate as a feedstock for *G. oxydans* only reduced bioleaching by 17% relative to glucose (this difference in performance is statistically significant; *p* < 0.001).

As cellulosic hydrolysate is mildly acidic (pH < 6), it appears that it could be more effective at bioleaching dunite than glucose (**Figure 9**). However, statistical significance tests from this experiment indicated no significant difference between sterile glucose and cellulosic hydrolysate controls (*p* = 0.821). Glucose and cellulosic hydrolysate feedstocks for bioleaching by *S. desiccabilis* were not significantly different (*p* = 1.000) (**Figure 9**). Cellulosic hydrolysate was about 16% more effective as a bioleaching feedstock than glucose with *P. simplicissimum*; but this result lacked statistical significance (*p* = 0.947). Results of statistical significance tests are shown in **Table S3**.

### *G. oxydans* Might Oxidize Iron

In this test and in several other experiments, we observed that the dunite leachate produced by direct contact with *G. oxydans* turned a dark reddish brown (**Figure S1C**). The leachate visually resembled rust, and its color darkened over time. In contrast, when the biolixiviant was filtered and then applied to minerals (indirect leaching), the color of the leachate was gray to light green, as expected from dissolved olivine. In contrast to *G. oxydans*, the leachate produced by both direct and indirect leaching with *P. simplicissimum* was a light green in all tests up to even 20 days.

The leachate color change observed when *G. oxydans* contacted minerals directly suggests that it could oxidize iron. This possible iron oxidation requires further investigation. If iron can serve as an electron source for *G. oxydans* after reduced carbon sources are nearly depleted, for example, this would imply that reduced iron may enhance *G. oxydans*’ survival and perhaps even its function. For biomining Fe^2+^-rich ultramafic materials, an iron-oxidizing microbe may then use more of the available reduced carbon for organic acid production and assimilation than as an energy source. This could enhance bioleaching performance and decrease resource requirements and associated costs. Alternatively, *G. oxydans* may oxidize Fe^2+^ to Fe^3+^ to decrease Fe^2+^ toxicity^65^, whereas *P. simplicissimum* may experience Fe^2+^ toxicity if it cannot oxidize Fe^2+^. Whether for energy use or to decrease toxicity, *G. oxydans* is not known to be capable of iron oxidation from the authors’ literature review.

### Bioleaching Smooths Dunite Particles’ Surfaces By Dissolution

Crystal fragments of olivine (∼ 30 - 70 µm) from the dunite were observed via SEM imaging to be coated in numerous fine-grained particles (∼ 1 - 2 µm) at the beginning of leaching experiments. In the sterile controls the olivine surfaces remained coated in fine grained particles of dunite after 5 days. Conversely, by day 5 of bioleaching many of the fine-grained particles had been dissolved, leaving behind a smoother olivine surface (**Figure 10**). Fine-grained particles have a greater surface area to volume ratio and thus dissolve more readily when exposed to the biolixiviant than larger particles. However, there is also evidence of dissolution of the larger dunite particles (especially when exposed to *G. oxydans*) with etch pits visible in SEM images by day 5 of bioleaching (**Figure 10E**). Notably, *G. oxydans* appears to adhere to the surface of olivine crystals which may further promote dissolution (**Figure 10 C and D**).

**Figure 10.**
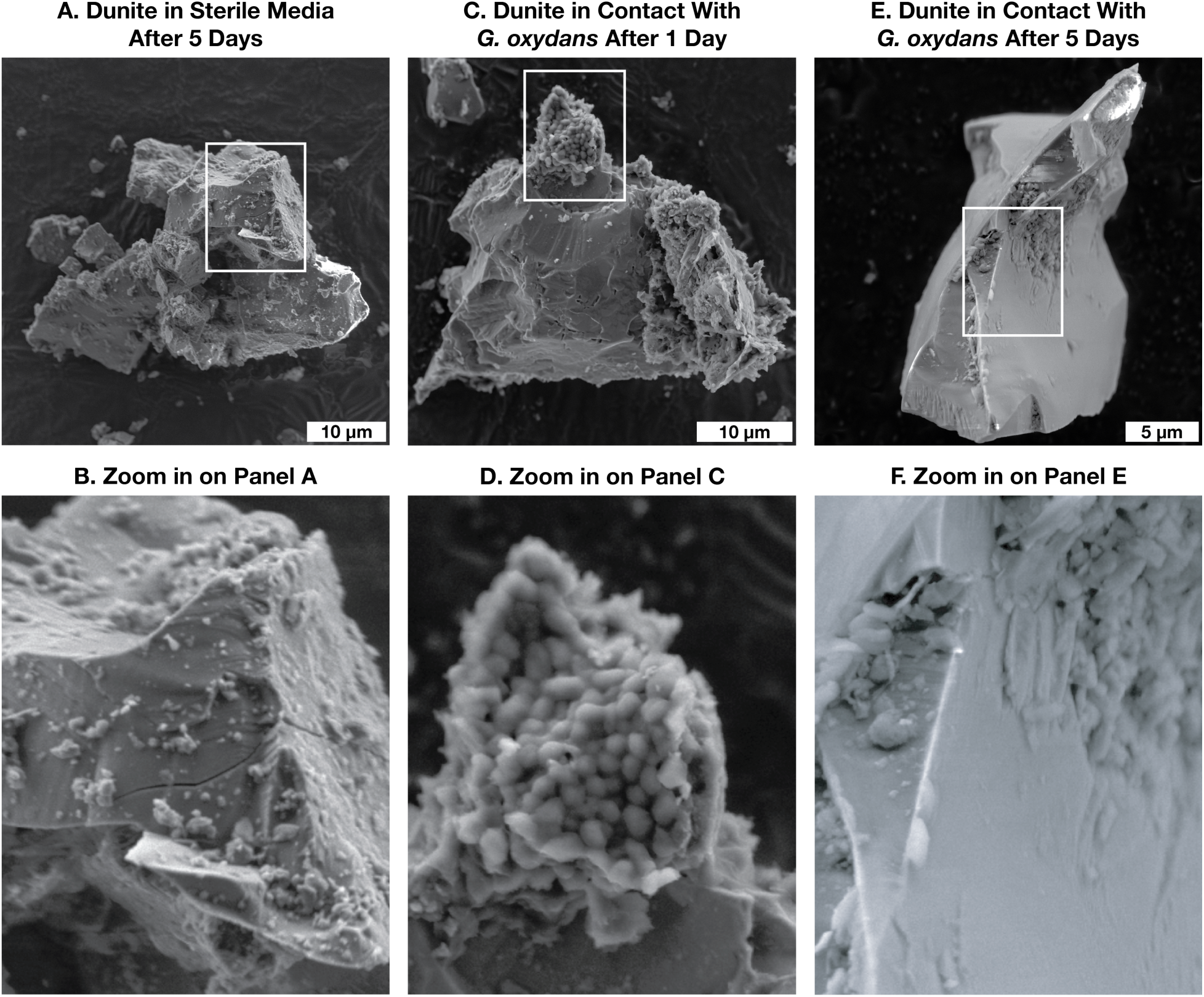
Bioleaching smooths the surfaces of dunite particles. (**A** and **B**) Dunite particles in sterile glucose controls display surfaces covered in fine-grained particles even after 5 days of leaching. (**C** and **D**) Dunite particles appear rough in texture after 1 day of leaching by *G. oxydans*. Notably, *G*. *oxydans* cells appear to adhere to the particles. (**E** and **F**) However, after 5 days of contact with *G. oxydans*, olivine crystals are noticeably smoothed, and most fine-grained particles have dissolved. Etch pits are also observed at the bottom of olivine particle in **(E).**

## Conclusion

The goal of this study was to find the most promising microbe out of three known candidates around which to build an accelerated weathering process for ultramafic rocks, and to further genetically engineer. Based on our results, *G. oxydans* is the clear choice of the three species. First, we have already demonstrated that its bioleaching capability can be improved by genetic engineering^9^. Second, out of the three microbes tested, here it is by far the best at leaching magnesium and energy-critical metals from ultramafic rock (**Figures 5** and **6**). Third, bioleaching by *G. oxydans* suffers far less inhibition at high pulp densities of ultramafic rock: at 2% pulp density, it is 117% better than its nearest competitor, *P. simplicissimum*; at 3% it is 398% better, and at 10% pulp density it is 1,713% better. This fact is of particular importance, as optimizing pulp density, particularly to allow for the use of high pulp densities, is an important factor in developing a viable industrial process. Fourth, *G. oxydans* can already use cellulosic hydrolysate as an effective substitute for glucose as a bioleaching feedstock (**Figure 9**).

However, this work is only the beginning of investigation into accelerated weathering of ultramafic materials. First, we need to investigate how bioleaching performance changes at much longer time durations than we investigated here (weeks to months). Second, further investigation is warranted for optimizing an engineered microbe for use with cellulosic hydrolysate, a cellulosic hydrolysate formula, and production protocol best suited for ultramafic bioleaching, and industrial bioleaching processes. We need to find out if we can engineer *G. oxydans* to be more tolerant of cellulosic hydrolysate and produce a biolixiviant that is equally, if not more, effective than the one derived from glucose. Third, we need to understand the mechanisms that *G. oxydans* uses for mineral-dissolution much better. We need to investigate if *G. oxydans* is capable of oxidizing iron, and if this contributes to its ability to tolerate high pulp densities of dunite, and if this can be improved. Finally, we need to understand how far the performance of ultramafic bioleaching can be pushed, and if it can be harnessed to improve the economics of bioleaching magnesium for CO_2_ sequestration and energy-critical elements needed for our future energy infrastructure.

## Supporting information

Dataset S1 - Ultramafic Rock Composition and Sequestration Potential

Dataset S2 - ICP-MS Data

## End Notes

### Data Availability

Raw ICP-MS data is included as **Dataset S2** with this article. Data for figures have been deposited on GitHub at https://github.com/barstowlab/article-044-three-microbes and archived on Zenodo^69^.

### Code Availability

No code was generated in the preparation of this article.

### Materials and Correspondence

Correspondence and material requests should be addressed to B.B.

## Acknowledgments

This work was made possible by support from a gift from Nancy and Bob Selander to B.B. and E.G.; the U.S. Army Advanced Civil Schooling Program; S.M. was supported by a Link Foundation Graduate Fellowship; J.A. was supported by an award from the Cornell Atkinson Center for Sustainability Academic Venture Fund. The cellulosic hydrolysate was provided as a generous gift from REPurpose (Horizon Europe program) from the Bio Base Europe Plant in Belgium. Correspondence with Rosa Santomartino made experiments with *P. simplicissimum* possible, and correspondence with Charles Cockell made experiments with *S. desiccabilis* possible. Stephen Parry of the Cornell Statistical Consulting Unit provided generous support in statistical analysis. Views and opinions expressed or implied herein are solely those of the authors and should not be construed as policy or carrying the official sanction of the Department of Defense, United States Army, or other agencies or departments of the U.S. Government.

## Contributions

Conceptualization, L.P., E.G., and B.B.; Methodology, L.P., J.D.K., J.L., A.H., J.A., S.M., M.C.R., E.G., and B.B.; Investigation, L.P., J.D.K., J.L., A.H.; Writing—Original draft, L.P.; Writing—Review and editing, L.P., J.D.K., E.G., and B.B.; Funding acquisition, E.G., and B.B.; Resources, M.C.R., E.G., and B.B.; Supervision, E.G. and B.B.; Data curation, L.P. and B.B.; Visualization, L.P. and B.B.; Formal analysis, L.P.

## Materials and Methods

### Media and Microbial Cultures

Yeast extract, peptone, and mannitol media (YPM) was prepared using methods DSMZ described previously^70^. Reasoner’s 2A media (R2A) was prepared using methods Reasoner described previously^71^. Yeast extract, peptone, and dextrose media (YPD) was prepared using methods Cold Spring Harbor Protocols described previously^72^.

The cellulosic hydrolysate source was REPurpose (Horizon Europe program) from the Bio Base Europe Plant in Belgium. The cellulosic hydrolysate was made from used cardboard. Initial concentrations of organic carbon-containing molecules were determined to be 562.44 g/L glucose, 101.34 g/L xylose, 32.04 g/L lactate, and 1.18 g/L acetate using high-performance liquid chromatography (HPLC). The liquid was diluted to 40% glucose weight / volume, filtered with 0.22 μm filters (Corning product number 430513), and stored at -20 ^°^C until needed.

#### Gluconobacter oxydans

The culture source was the American Type Culture Collection (ATCC) as type strain B58. It was and was grown on YPM plates at room temperature, frozen at -80 ^°^C in 30% glycerol, and then cultured for three days on YPM plates at room temperature. Plates were stored at 4 ^°^C and used within two weeks.

#### Sphingomonas desiccabilis

The culture source the American Type Culture Collection (ATCC) as type strain CP1D DSM 16792. It was grown on R2A plates at room temperature, frozen at -80 ^°^C in 30% glycerol, and then cultured for two days on YPM plates at room temperature. Plates were stored at 4 ^°^C and used within two weeks.

#### Penicillium simplicissimum

The culture source was the German Collection of Microorganisms and Cell Cultures GmbH freeze type strain DSMZ 1078. It was grown on YPD plates at room temperature, frozen at -80 ^°^C in 30% glycerol, and then cultured for five days on YPM plates at room temperature. The mycelium was very gently scratched and rinsed with 0.9% sterile saline solution. Spore counts were determined to be ∼10^6^ using an Improved Neubauer hemocytometer. Spore solutions were then stored at 4 ^°^C and used within two weeks.

Optical densities (OD) were read at 590 nm using a WPA Colourwave CO7000. All pHs were read using a VWR Symphony B10P. Erlenmeyer flasks used were VWR 100 mL Cat. No. 75804-644. Holes were bored into #7 neoprene stoppers with a 3/16” drill bit, stoppers were washed, and 0.2 μm hydrophobic filters (Pall product number 4225T) were inserted into the tops of the stoppers. These filters allowed aseptic gas exchange while preventing evaporation.

#### Day Book Dunite

Ultramafic samples were collected from the Day Book dunite deposit in western North Carolina^73^. The chemically weathered exterior of the dunite was removed using an MK Diamond tile saw (model MK-101). The unaltered interior was pulverized into powder using a SPEX SamplePrep 8000M mill and tungsten carbide vial. The mineralogy of the powder before bioleaching was characterized via x-ray diffraction using a Bruker D8 Advance ECO powder diffractometer. The dunite is composed of ∼92 wt. % olivine (Mg_2_SiO_4_), 5 wt. % antigorite ((Mg,Fe)_3_Si_2_O_5_(OH)_4_), 2 wt. % lizardite (Mg_3_Si_2_O_5_(OH)_4_), and 1 wt. % talc (Mg_3_Si_4_O_10_(OH)_2_). The bulk rock major and trace element compositions were measured at Hamilton Analytical Laboratories via x-ray fluorescence (XRF; Thermo ARL Perform’X spectrometer) and inductively-coupled plasma mass spectrometry (ICP-MS; Agilent 7800 Mass Spectrometer at Cornell CMaS facility), respectively. Following leaching, dunite experimental residues were imaged and chemically mapped using scanning electron microscope energy dispersive spectroscopy (SEM-EDS) to characterize alteration induced by the bioleaching. Finally, the mineralogy of residues was once again measured via x-ray diffraction to identify any new minerals produced during the bioleaching experiments.

The powdered dunite was sterilized at 135 ^°^C and 30 pounds per square inch (psi; ≈ 200 megapascals) over atmospheric pressure for three minutes in an autoclave. The dunite was then weighed to within 1% of the required weight for the applicable pulp density and liquid volume.

### Work Flow for Three Microbes’ Baseline Performance at 1%, 2%, and 3% Pulp Densities

Liquid cultures were started for the two bacteria in 25 mL YPM in 100 mL VWR flasks. Technical replicates were used in this experiment: a single *G. oxydans* colony inoculated a single flask of 25 mL YPM, and a single *S. desiccabilis* colony inoculated a single flask of 25 mL YPM for each species for each pulp density experiment. These liquid cultures were grown at 30 ^°^C for two days with mixing at 200 rotations per minute (rpm). Three total experiments occurred – one each for 1% dunite, 2% dunite, and 3% dunite pulp densities (solid to liquid ratios). Initial inoculation of flasks included back-diluting the single flasks for each bacterium into 18 flasks per bacterium to 0.05 OD in 10 mL YPM. The 18 flasks per bacterium included triplicates for sample days 0, 1, 2, 3, 4, and 5. Initial inoculation of the fungus ’18 flasks in each experiment included 1% (0.1 mL) of a ∼10^6^ spores solution in 10 mL YPM. These flasks for all species were grown at 30 ^°^C for two days with no mixing. After each microbe grew for two days in the YPM growth medium, 5 mL of a 40% w/v glucose solution (400 g/L) were added to each flask for a total volume of 15 mL. Dunite was also added with the glucose. For the 1% pulp density experiment, 0.15 g dunite was added to each flask; 0.30 g was added for 2%, and 0.45 g was added for 3%. Thus, normalization for comparing the very different species included using consistent time, media fed, and growing conditions among all microbes for all tests.

Samples were collected by pouring them into 50 mL centrifuge tubes and centrifuging them at 3,214 × *g* for 10 minutes. After this, 1.5 mL of the leachate was stored at -80 ^°^C for future ICP-MS analysis. Then, pH was read for the remaining leachate. This leachate was then discarded, and the leached rocks and microbes were rinsed with DI water and dried at room temperature for future analysis. Fungus growth, however, was removed using inoculation picks prior to centrifugation. Fungal growth floated on the surface of the liquid in the flasks, whereas the bacterial growth was mixed with the powdered minerals on the bottoms of the flasks (**Figure S1**).

Sterile controls consisted of 13.3% w/v glucose and applicable dunite for each pulp density. Controls were also set in triplicate per sample day for a total of 18 flasks per each of the three pulp density experiments.

### Work Flow for Cellulosic Hydrolysate Experiment

Tests comparing the three microbes’ performance using glucose versus cellulosic hydrolysate as feedstocks occurred with 15 days of leaching at room temperature, with 10% dunite pulp density, and with flask liquid volumes of 10 mL. Biological replicates were used for the different bacteria: a colony inoculated 3 mL of YPM in a culture tube grown at two days in 30 ^°^C at 200 rpm, then the tube growth was diluted into a flask of 10 mL YPM to 0.05 OD, then the flask was grown at 30 ^°^C at 200 rpm for 2 days, the culture was mixed and reduced in volume to 5 mL by discarding mixed culture, and 5 mL 40% w/v glucose or 40% w/v cellulosic hydrolysate-glucose and 1 g dunite were added.

Each microbe had six flasks: three for glucose and three for cellulosic hydrolysate, so six colonies were used for each species. For the fungus, 1 mL of spores in YPM was added to 4 mL YPM, 5 mL 40% w/v glucose or cellulosic hydrolysate-glucose, and 1g dunite. Flasks were then set at room temperature with stoppers preventing air exchange and evaporation, and further growth and bioleaching occurred for 15 days.

Sterile controls were set in triplicate with 20% w/v glucose or cellulosic hydrolysate-glucose and 1g dunite each for a total of six controls.

### ICP-MS

Frozen samples were thawed, centrifuged in 0.45 µm polypropylene 96-well filter plates into polypropylene 96 well-plates for 10 min at 3,214 x *g*, and they were then diluted to 1% of their original concentration in 5 mL of a 2% nitric acid solution. Next, the samples were analyzed by an Agilent 7800 inductively coupled plasma-mass spectrometry (ICP-MS). The ICP-MS standard was custom-made by High Purity Standards at 1,000 µg/mL iron, magnesium, and silicon and 100 µg/mL calcium, cobalt, chromium, copper, potassium, manganese, sodium, nickel, phosphorus, titanium, and zinc.

### Statistical Analysis

A one-way ANOVA test compared all leachate magnesium concentrations in the glucose versus cellulosic hydrolysate experiment (**Table S3A**). This was followed by a Turkey test to compare each species’ leachate magnesium concentrations using different sugar sources (**Table S3B**). Jamovi was used for these statistical significance tests^74^.

