## Supplementary Information for "Cross-species Comparison of Ultramafic Rock Bio-accelerated Weathering Performance"

### Supplementary Information Figures

**Figure S1.** Set up of bioleaching experiments.

### Supplementary Information Tables

**Table S1**. Dunite leachate minimum pH summary.

**Table S2**. Dunite leachate maximum $Mg^{2+}$ and $Ni^{2+}$ concentration summary.

**Table S3.** Statistical significance tests on dunite bioleaching with cellulosic hydrolysate and glucose feedstocks.

### Supplementary Information Datasets

**Dataset S1.** Ultramafic rock composition and $CO_2$ sequestration potential.

**Dataset S2.** Mass spectrometry data used in this article.

### Supplementary Information Notes

**Note S1.** Total $CO_2$ addition to the atmosphere since the start of the Industrial Revolution.

**Note S2.** Estimates of time for draw down of anthropogenic $CO_2$ by natural processes.

**Note S3.** Estimates for $CO_2$ sequestration potential of surface-accessible ultramafic material.

### Supplementary Figures

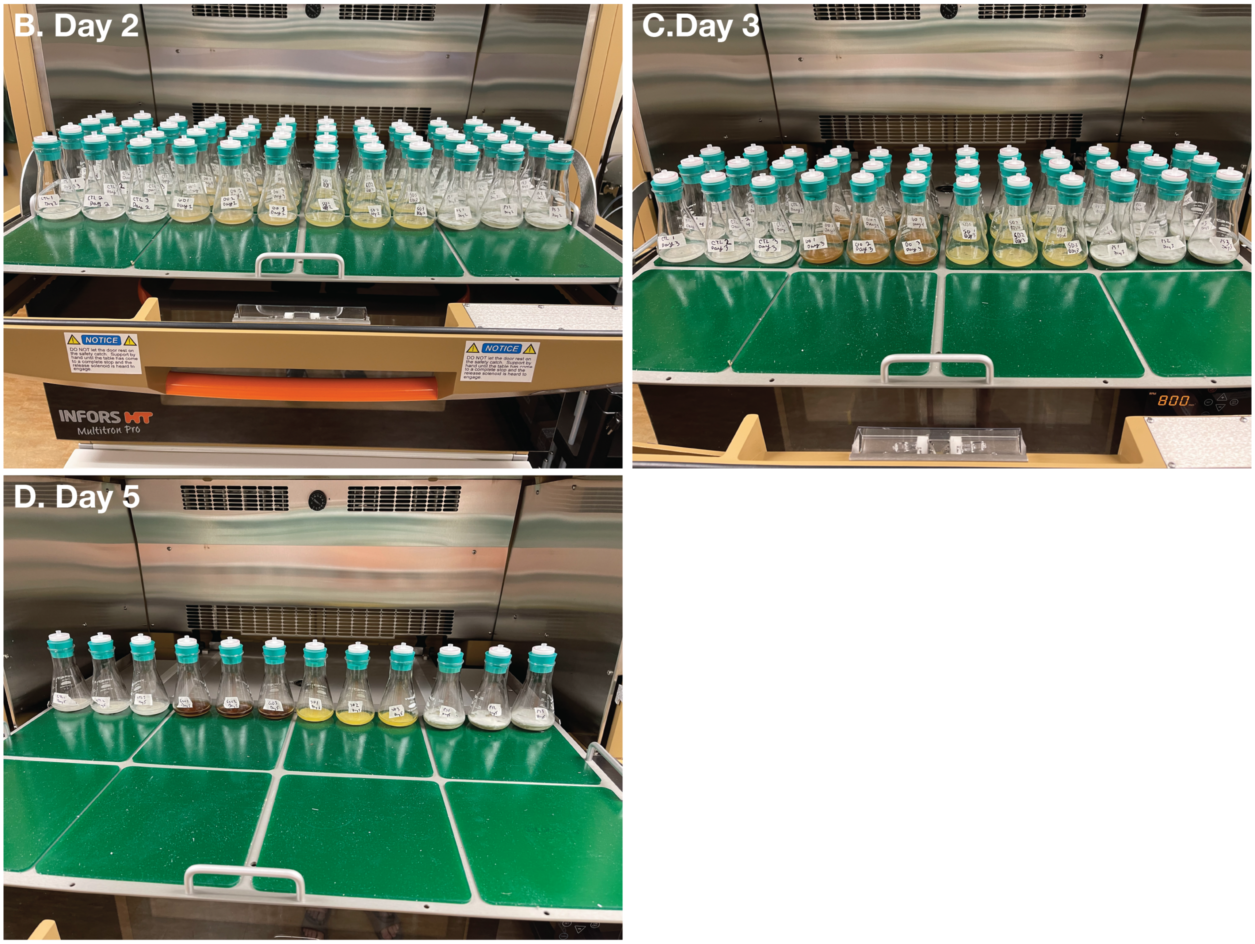


**Figure S1.** Set up of bioleaching experiments. At the start of the experiment, flasks were set up identically, with some flasks pre-scheduled for removal. For example, some flasks were intended for removal on day 1, while others were not intended to be removed until day 5. (**A**) Day 2 of bioleaching. Note that the flasks in the three columns furthest to the right contain the floating fungus *Penicillium simplicissimum*. (**B**) Day 3 of bioleaching. (**C**) Day 5 of bioleaching. Note the $4^{th}$, $5^{th}$, and $6^{th}$ flasks from the left containing *G. oxydans* (Go) that show heavy oxidation.

### Supplementary Tables

| Organism | Minimum pH | Day Observed | Pulp Density (%) |
| --- | --- | --- | --- |
| *G. oxydans* | 2.65 ± 0.032 | 3 | 3 |
| *P. simplicissimum* | 4.44 ± 0.031 | 4 | 3 |
| *S. desiccabilis* | 6.68 ± 0.020 | 5 | 3 |
| Sterile control | 8.40 ± 0.025 | 0 | 3 |
| *G. oxydans* | 2.54 ± 0.031 | 3 | 2 |
| *P. simplicissimum* | 4.08 ± 0.036 | 5 | 2 |
| *S. desiccabilis* | 5.95 ± 0.032 | 5 | 2 |
| Sterile control | 8.31 ± 0.006 | 4 | 2 |
| *G. oxydans* | 2.15 ± 0.057 | 3 | 1 |
| *P. simplicissimum* | 3.75 ± 0.032 | 4 | 1 |
| *S. desiccabilis* | 5.53 ± 0.080 | 5 | 1 |
| Sterile control | 8.12 ± 0.049 | 0 | 1 |

**Table S1. Dunite leachate minimum pH summary.** This table summarizes key results of bioleaching of dunite from **Figures 1** and **2** in the main text. The sterile control contained 13.3% w/v glucose.

| Organism | Maximum $Mg^{2+}$ Concentration (mg/L) | Day Observed | Maximum $Ni^{2+}$ Concentration (mg/L) | Day Observed | Pulp Density (%) |
|---|---|---|---|---|---|
| *G. oxydans* | 1,064 ± 83 | 5 | 11.0 ± 0.60 | 5 | 1 |
| *P. simplicissimum* | 487 ± 26 | 5 | 5.65 ± 0.22 | 5 | 1 |
| *S. desiccabilis* | 65 ± 22 | 5 | 1.21 ± 0.24 | 5 | 1 |
| Sterile control | 24 ± 0.58 | 3 | 0.72 ± 0.056 | 1 | 1 |
| *G. oxydans* | 1,973 ± 49 | 5 | 19.5 ± 0.67 | 5 | 2 |
| *P. simplicissimum* | 907 ± 15 | 5 | 9.99 ± 0.076 | 5 | 2 |
| *S. desiccabilis* | 73 ± 1.9 | 5 | 1.49 ± 0.027 | 5 | 2 |
| Sterile control | 27 ± 1.3 | 4 | 0.547 ± 0.010 | 2 | 2 |
| *G. oxydans* | 3,130 ± 64 | 5 | 33 ± 0.62 | 5 | 3 |
| *P. simplicissimum* | 629 ± 45 | 5 | 7.62 ± 0.36 | 5 | 3 |
| *S. desiccabilis* | 99 ± 3.1 | 5 | 2.08 ± 0.085 | 5 | 3 |
| Sterile control | 59 ± 0.75 | 4 | 1.91 ± 0.14 | 2 | 3 |

**Table 2. Dunite leachate maximum $Mg^{2+}$ and $Ni^{2+}$ concentration summary.** This table summarizes key results of bioleaching of dunite from **Figures 3**, **4**, and **6** in the main text. The sterile control contained 13.3% w/v glucose.

*A. Welch's One Way ANOVA Test*

| | F | df1 | df2 | $p$-value |
|---|---|---|---|---|
| **$Mg^{2+}$ Bioleaching (mg/L)** | 1015 | 7 | 6.63 | < 0.001 |

*B. Tukey's Post-Hoc Test*

| Sample | | Ctrl CH | Ctrl G | *Go* CH | *Go* G | *Ps* CH | *Ps* G | *Sd* CH | *Sd* G |
|---|---|---|---|---|---|---|---|---|---|
| **Ctrl CH** | Mean diff. | - | 307 | -6921 | -8408 | -1280 | -1046 | -31.1 | -93.5 |
| | $p$-value | - | 0.821 | < .001 | < .001 | < .001 | 0.003 | 1.000 | 1.000 |
| **Ctrl G** | Mean diff. | | - | -7228 | -8715 | -1587 | -1353 | -338.4 | -400.9 |
| | $p$-value | | - | < .001 | < .001 | < .001 | < .001 | 0.745 | 0.573 |
| ***Go* CH** | Mean diff. | | | - | -1487 | 5641 | 5875 | 6889.8 | 6827.4 |
| | $p$-value | | | - | < .001 | < .001 | < .001 | < .001 | < .001 |
| ***Go* G** | Mean diff. | | | | - | 7128 | 7362 | 8376.9 | 8314.5 |
| | $p$-value | | | | - | < .001 | < .001 | < .001 | < .001 |
| ***Ps* CH** | Mean diff. | | | | | - | 234 | 1248.8 | 1186.4 |
| | $p$-value | | | | | - | 0.947 | < .001 | < .001 |
| ***Ps* G** | Mean diff. | | | | | | - | 1015.0 | 952.6 |
| | $p$-value | | | | | | - | 0.004 | 0.007 |
| ***Sd* CH** | Mean diff. | | | | | | | - | -62.4 |
| | $p$-value | | | | | | | - | 1.000 |
| ***Sd* G** | Mean diff. | | | | | | | | - |
| | $p$-value | | | | | | | | - |

**Table S3.** Statistical significance tests on dunite bioleaching with cellulosic hydrolysate (CH) and glucose (G) feedstocks. (**A**) Comparison of all bioleaching test species and conditions to each other by a one-way ANOVA test indicates statistical significance of differences in performance. (**B**) Statistical significance comparisons between entries in **Figure 8** in the main text. Ctrl: Sterile control; *Go*: *Gluconobacter oxydans*; *Ps*: *Penicillium simplicissimum*; *Sd*: *Sphingomonas desiccabilis*. Calculations were performed with jamovi [jamovi2022a].

### Supplementary Notes

#### Note S1. Total $CO_2$ Addition to the Biosphere Since the Start of the Industrial Revolution

Given the precision with which atmospheric $CO_2$ concentrations can be measured (either through direct measurement or by ice core data), there is a surprisingly high degree of uncertainty on the amount of $CO_2$ that has been added to the biosphere by burning of fossil fuels since the start of the Industrial Revolution (approximately 1750).

Estimates for the total $CO_2$ addition to the biosphere range from a low estimate of approximately 1 trillion tonnes [NOAA2021a], to an often-quoted estimate of 1.5 trillion tonnes [Keller2018a, Lackner2021a], with a recent high estimate of 1.73 trillion tonnes [Andrew2023a]. Assuming that surface-accessible rocks can only sequester 100 trillion tonnes of $CO_2$ as carbonate minerals [Kelemen2019a], this is still 58 times more than the highest estimate of excess $CO_2$ in the atmosphere [Andrew2023a].

#### Note S2. Estimates of Time for Draw Down of Anthropogenic $CO_2$ by Natural Processes.

As with estimates of the total $CO_2$ addition to the biosphere (**Note S1**), there is also a surprising amount of uncertainty on the time it will take for natural processes to remove anthropogenic $CO_2$ from the biosphere.

Gaillardet *et al*. [Gaillardet1999a] estimated that the magnesium and calcium cations available in rivers in streams could remove $CO_2$ from the atmosphere at a rate of 381 million tonnes per year. More recently, Hartmann *et al*. [Hartmann2009a] estimated that weathering and subsequent carbon mineralization removes 871 million tonnes to 1.06 billion tonnes of $CO_2$ annually. Hartmann's estimate of $CO_2$ removal suggests that Gaillardet's estimate does not account for all $CO_2$ mineralization processes. Recently, Zondervan *et al*. estimated the release of 249 million tonnes of $CO_2$ per year from oxidation of rock organic carbon [Zondervan2023a], that offsets carbon mineralization. However, both Gaillardet's and Hartmann's estimates of total $CO_2$ mineralization exceed Zondervan's estimate of $CO_2$ release.

By subtracting Zondervan's estimate of $CO_2$ release from the $CO_2$ removal rates via mineralization estimated by Hartmann we can estimate a net $CO_2$ removal rate of approximately 622 to 811 million tonnes of $CO_2$ per year. Assuming a linear draw down of $CO_2$, the time to remove anthropogenic $CO_2$ is approximately 2,130 to 2,780 years (assuming 1.73 trillion tonnes of anthropogenic $CO_2$). However, evidence from the geological record suggests that after rapid warming periods (*e.g.*, the Paleocene-Eocene Thermal Maximum), the time needed to draw down excess $CO_2$ was much longer than a simple linear model suggests. As a result, removal of excess $CO_2$ from the atmosphere caused by anthropogenic emissions could extend into tens or even hundreds of thousands of years [Rohl2007a]

### Note S3. Estimates for $CO_2$ Sequestration Potential of Surface-Accessible Ultramafic Material.

Carbon mineralization proceeds through a two step process. In the first, ultramafic material is dissolved by weathering to release magnesium. In a second step, magnesium is reacted with dissolved carbonate to form magnesite [Power2013a],

$$Mg^{2+}_{aq} + CO^{2-}_{3,\,aq} \rightarrow MgCO_{3,\,s}. \tag{S1}$$

How much ultramafic material needs to be dissolved to capture 1 tonne of $CO_2$? Comparing surveys representative of ultramafic rock composition (Oman Peridotite, Josephine Ophiolite, Abyssal Peridotite, and Cratonic Peridotite), and we converted these values from the standard oxide weights to percent of metals by weight. Results from this conversion show that ultramafic rocks, on average, are comprised of about 26.5 % Mg by weight [Hanghoj2010a, LeRoux2014a, Niu2004a, Wittig2008a] (**Dataset S1**),

$$f_{\mathrm{Mg}} = 0.2654. \tag{S2}$$

Meanwhile, the ratio of molecular weights of magnesium and $CO_2$,

$$\begin{aligned} r_{\mathrm{C-Mg}} &= \mathrm{MW}_{\mathrm{CO_2}} / \mathrm{MW}_{\mathrm{Mg}}, \\ &= 44.009/24.305, \\ &= 1.811. \end{aligned} \tag{S3}$$

Thus, assuming one mol of magnesium sequesters one mol of $CO_2$, a given mass, $M_{\mathrm{ultramafic}}$ of ultramafic material can sequester $M_{\mathrm{seq}}$ of $CO_2$,

$$M_{\mathrm{seq}} = M_{\mathrm{ultramafic}}\, r_{\mathrm{C-Mg}}\, f_{\mathrm{MgO}}. \tag{S1}$$

Therefore, 1 tonne of ultramafic rock can sequester 0.4806 tonnes of $CO_2$ (**Dataset S1**). This estimate is consistent with previous estimates of individual ultramafic rock sequestration potentials [NAS2019a]. Therefore, 400 megatonnes of ultramafic material (annual mine tailings production [Power2013a]) can sequester 192 megatonnes of $CO_2$. Meanwhile, 10 gigatonnes of ultramafic material (best estimate of global stockpile of mine tailings [Power2013a]) can sequester 4.806 gigatonnes of $CO_2$.
